# Autoimmune diabetes-dependent c-Maf SUMOylation licenses inflammatory bowel disease by reshaping the gut microbiota

**DOI:** 10.64898/2026.06.15.732285

**Authors:** Chao-Yuan Hsu, Yi-Wen Tsai, Shin-Huei Fu, Yu-Wen Liu, Jia-Ling Dong, Yun-Ju Yang, Yu-Wen Mai, Ling-Chun Tsai, Chi-En Wu, Hsin-I Liang, Chi-Chin Sun, Chien-Tzung Chen, Shu-Ping Wang, Shi-Chuen Miaw, Huey-Kang Sytwu

**Affiliations:** Graduate Institute of Microbiology and Immunology, College of Biomedical Sciences, National Defense Medical University, Taipei City, Taiwan; Graduate Institute of Life Sciences, College of Biomedical Sciences, National Defense Medical University, Taipei City, Taiwan; Graduate Institute of Medical Sciences, College of Medicine, National Defense Medical University, Taipei City, Taiwan; Department of Family Medicine, Chang Gung Memorial Hospital, Linkou, Taoyuan City, Taiwan; School of Medicine, College of Medicine, Chang-Gung University, Taoyuan City, Taiwan; National Institute of Infectious Diseases and Vaccinology, National Health Research Institutes, Miaoli County, Taiwan; Department of Neurological Surgery, Tri-Service General Hospital, School of Medicine, National Defense Medical Center, Taipei, Taiwan; Department of Obstetrics and Gynecology, Taichung Armed Forces General Hospital, Taichung City, Taiwan; Department of Ophthalmology, Chang Gung Memorial Hospital, Keelung, Keelung City, Taiwan; Department of Plastic and Reconstruction Surgery, Chang Gung Memorial Hospital, Linkou, Taoyuan City, Taiwan; Institute of Biomedical Sciences, Academia Sinica, Taipei City, Taiwan; Graduate Institute of Immunology, National Taiwan University College of Medicine, Taipei City, Taiwan

**Keywords:** Type 1 diabetes, Inflammatory bowel disease, c-Maf SUMOylation, IL-21, *Lactobacillus johnsonii*, Lithocholic acid and HDAC2

## Abstract

We have previously demonstrated a critical role of c-Maf SUMOylation in the regulation of autoimmune diabetogenesis, but its physiological relevance to and potential clinical impact on gut inflammation need further elucidation. Here, integrating a 14-year population-based time-trend cohort study of 139,204 type 1 diabetes patients with experiments in non-obese diabetic mice, we illustrated that autoimmune diabetes confers resistance to colitis mediated by an impaired c-Maf SUMOylation-driven IL-21-IgA axis. Utilizing T cell-specific c-Maf SUMOylation site-mutated mice, we further demonstrated that SUMOylation-defective c-Maf enhances IL-21 expression in CD4^+^ T cells to promote fecal IgA production and colitis resistance via microbiota remodeling, specifically through *Lactobacillus johnsonii* enrichment and activating lithocholic acid (LCA)-mediated AMPK anti-inflammatory pathway. Pharmacological HDAC2 inhibition by BRD6688 promotes c-Maf-mediated IL-21 and suppresses colitis in PBMC-humanized mice. Altogether, we revealed how SUMOylation reciprocally modulates the inflammatory process between autoimmune diabetes and colitis in a T cell-restricted and single transcription factor-based manner.

## Introduction

Inflammatory bowel disease (IBD) comprises ulcerative colitis (UC) and Crohn’s disease, and is a relapsing–remitting disorder of the gastrointestinal tract that affects millions globally and imposes a substantial burden on health systems^1,2,3^. Despite the advent of biological agents and immunomodulatory therapies, current treatments largely alleviate the symptoms rather than prevent disease progression, and many patients experience progressive mucosal damage and eventual loss of therapeutic responsiveness^4,5,6^. Although genome-wide association studies have identified numerous loci associated with IBD susceptibility, the genetic contribution alone is insufficient to explain the observed clinical heterogeneity or the escalating global incidence^7,8^. These limitations highlight an urgent need for mechanism-based interventions that can restore durable intestinal immune tolerance and reestablish mucosal homeostasis. The multifaceted pathogenic network involving immune imbalance and environmental stressors, particularly microbial dysbiosis, has been implicated in driving the development and persistence of IBD^9,10^. Preclinical models, such as dextran sulfate sodium (DSS)-induced colitis, have demonstrated that inflammatory severity is not determined solely by the host genotype but is also affected by the interplay between IgA-dependent immunological signals and the microbial profile within the gut environment^11,12,13^. However, the precise molecular mechanisms that confer resilience and/or promote inflammatory escalation remain poorly characterized.

The genetic and epidemiological links between IBD and T1D are well documented^14^. Current studies have reported a positive correlation between these 2 diseases, which suggests that genetic and environmental factors contribute to this association^15,16,17^. However, additional genetic studies have suggested paradoxically that T1D-associated loci may confer protection against IBD development^18,19^. The discrepancies between observational studies highlight the inherent limitations and challenges in drawing causal inferences from real-world data, and these studies are often affected by bias and nonrandomized exposure patterns. Few studies have integrated population-based findings with experimental dissection of the underlying immunological mechanisms. In this study, we addressed this issue further to reconcile this discrepancy by combining large-scale and longitudinal epidemiological analysis with a mechanistic investigation of T-cell-intrinsic signaling pathways. Here, we report our findings that confirm an inverse correlation between IBD and T1D. In addition, we describe a causal immune–microbiota axis to explain how disease-dependent programming in T1D confers protection against intestinal inflammation.

SUMOylation critically modulates immune signaling networks and transcriptional programming, particularly within T cells^20,21,22,23,24^. Through the reversible covalent conjugation of small ubiquitin-like modifier (SUMO) proteins to specific lysine residues, this modification orchestrates a wide range of cellular processes, including transcription factor stabilization, nuclear–cytoplasmic trafficking, chromatin remodeling, and protein–protein interactions^25,26,27^. These molecular effects enable the precise control of gene expression programs that are fundamental to maintaining peripheral immune tolerance and suppressing unwarranted inflammatory responses^28,29,30,31^. Emerging studies have increasingly implicated dysregulation of the SUMOylation machinery in the initiation and progression of T1D and IBD^24,31^.

We have previously identified a key transcription factor c-Maf as a direct substrate of SUMOylation and have demonstrated that its SUMOylation limits the transcriptional activation of proinflammatory genes and restricts diabetogenic T-cell effector functions in NOD mice. These findings underscore the immunosuppressive role of SUMOylation in autoimmune diabetes^24^. However, it remains unclear whether this SUMOylation-based regulatory axis is implicated similarly in the context of mucosal immune responses and intestinal inflammation. Considering the tissue-specific microenvironmental cues and transcriptional plasticity of gut- resident T cells, we asked whether c-Maf SUMOylation alone is sufficient to modulate the diverse T-cell subsets implicated in IBD pathogenesis. If so, further studies would be warranted to elucidate whether targeting SUMOylation pathways in a T-cell-restricted and transcription factor-specific manner might provide a feasible strategy for mitigating intestinal inflammation. In this study, we conducted a comprehensive nationwide longitudinal cohort analysis and integrated population-scale epidemiological data with an in vivo mechanistic investigation using a T-cell-specific c-Maf SUMOylation-defective mouse model. This 2-pronged approach allowed us to dissect the functional consequences of impaired SUMOylation within a defined immune compartment. Through these complementary investigations, we demonstrate that targeted disruption of c-Maf SUMOylation in T cells potentiates an IL-21-driven IgA class switching, which then increases the mucosal immunoglobulin responses. This immunological shift significantly remodels the composition and diversity of the gut microbiota by favoring a microbial milieu associated with mucosal immune protection. Functionally, these c-Maf SUMOylation-defective mice exhibited stronger resistance to DSS-induced colitis compared with a control group, which suggests that the absence of c-Maf SUMOylation reconfigures T-cell-dependent regulatory networks and promotes intestinal resilience to inflammation. Collectively, these findings unveil a previously unappreciated immunoregulatory circuit in which posttranslational modification of a single transcription factor governs mucosal immunity through modulation of microbiota. This axis provides both mechanistic explanation into the inverse correlation between T1D and IBD and offers a conceptual framework in which targeting cell-specific SUMOylation may serve as a unified therapeutic strategy for balancing immune responses across distinct tissue environments.

## Results

### Autoimmune diabetes reduces susceptibility to IBD in both human studies and mouse models

Both genetic and epidemiological investigations have established a comorbid association between T1D and IBD, particularly in European populations^14,15,16^. However, contradictory studies have reported that certain alleles associated with T1D may have a protective effect against the development of IBD^18,19^, a finding that suggests a complex genetic landscape and underscores the need for further investigation of the mechanisms involved.

To address this discrepancy, we first conducted a retrospective population-based analysis using the National Health Insurance Research Database (NHIRD) in Taiwan, followed by a mouse-based mechanistic study. We identified 155,101 individuals diagnosed with T1D included in the NHIRD diabetes registry since January 1, 1997. After application of the exclusion criteria, 139,204 T1D patients and an equal number of matched nondiabetic controls were selected based on age, sex, and multiple comorbidities (Table 1). Both groups had a mean age of 57.57 years; the age distributions of the patient and control groups were as follows: 0–10 years (0.78% vs. 0.77%), 10–20 years (2.55% vs. 2.55%), 20–30 years (3.07% vs. 3.07%), and >30 years (93.61% vs. 93.61%). The percentage of male participants was similar in the 2 populations (52.19% vs. 52.22%). Most of the enrolled participants were aged >30 years (93.61%, *n* = 130,303 for each group). Hypertension and dyslipidemia were the most prevalent comorbidities in the T1D group, and hypertension and liver cirrhosis predominated among the controls (data not shown). The incidence of IBD was lower in the T1D cohort than in the control cohort (0.01%, *n* = 11 vs. 0.02%, *n* = 22) (Table 1). Our epidemiological findings along with genetic research on Asia population ^19^ provide support for an inverse correlation between these 2 diseases, which reflect divergent immunogenetic trajectories. This inverse relationship but differences in immunogenetic trajectories warrant further investigations to delineate the mechanisms underpinning the interplay between these diseases.

**Table 1.** Baseline characteristics and ulcerative colitis incidence in T1DM and matched non-T1DM cohorts. Demographic characteristics and the prevalence of ulcerative colitis were analyzed using a 14-year population-based cohort from the National Health Insurance Research Database (NHIRD) in Taiwan. Patients with Type 1 Diabetes Mellitus (T1DM, n = 139,204) were matched with non-T1DM controls (n = 139,204) at a 1:1 ratio based on age and gender. The incidence of ulcerative colitis is presented as both total cases and cases per 100,000 inhabitants. Data are presented as mean for continuous variables and n (%) for categorical variables.

| Characteristics | T1DM(n=139204) |  | non-T1DM(n=139204) |  |
| --- | --- | --- | --- | --- |
|  | N(Mean) | %(SD) | N(Mean) | %(SD) |
| **Age, years** | 57.57 | 16.15 | 57.57 | 16.15 |
| **Age group** |  |  |  |  |
| 0-10 years | 1085 | 0.78% | 1077 | 0.77% |
| 10-20 years | 3544 | 2.55% | 3552 | 2.55% |
| 20-30 years | 4272 | 3.07% | 4272 | 3.07% |
| >30 years | 130303 | 93.61% | 130303 | 93.61% |
| **Gender** |  |  |  |  |
| Male | 72656 | 52.19% | 72687 | 52.22% |
| Female | 66548 | 47.81% | 66517 | 47.78% |
| **ulcerative colitis** |  |  |  |  |
| total | 11 | 0.01% | 22 | 0.02% |
| per 100,000 inhabitants | 7.9 |  | 15.8 |  |

Analysis of the cumulative incidence during the follow-up from 2000 to 2013 revealed a significant reduction in the concurrence of IBD among individuals with T1D compared with matched nondiabetic controls (Fig. 1a, left panel). A stratified evaluation indicated that both risks of developing UC and Crohn’s disease were reduced in the T1D group and that the disease-specific cumulative curves displayed lower trajectories in the T1D group relative to the controls (Fig. 1a, middle and right panels, respectively). These quantitative data support our hypothesis that T1D confers a lower risk for IBD. The results of the epidemiological analyses showing a reduced incidence of IBD in the T1D cohort suggest a divergence between the processes of pancreatic autoimmunity and intestinal inflammation.

**Fig. 1.**
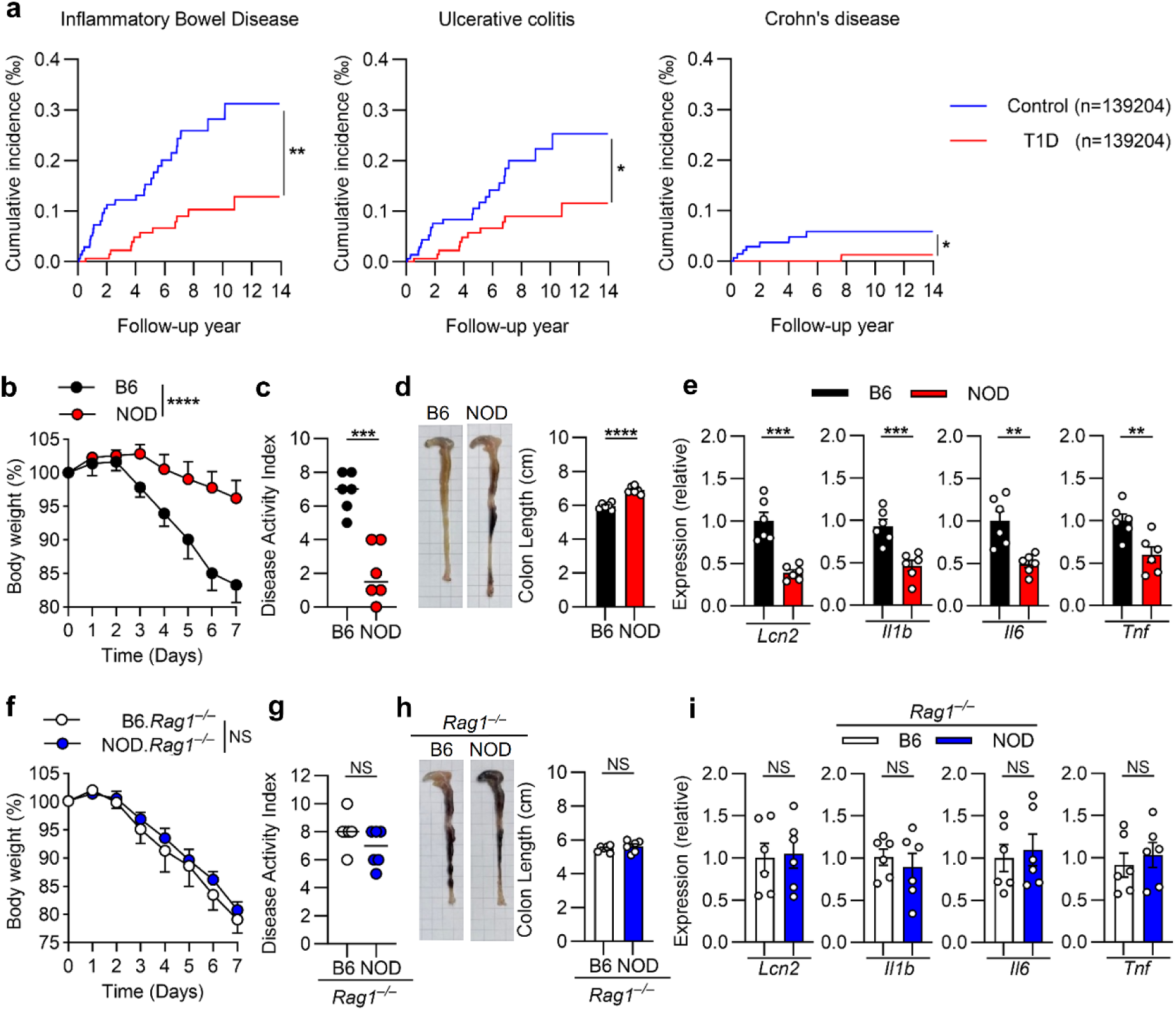
Autoimmune diabetes reduces susceptibility to inflammatory bowel disease in both human studies and mouse models. (**a**) The cumulative incidence of inflammatory bowel disease, ulcerative colitis and Crohn’s disease in patients with type 1 diabetes (T1D, red line, n=139,204) compared with age- and sex-matched non-T1D controls (Control, blue line, n = 139,204) over a 14-year follow-up period using a population-based cohort in Taiwan. (**b**–**e**) B6 and NOD mice were treated with 2.5% DSS for 7 days. Body weight changes (**b)** were monitored daily. Disease activity index (**c**), the colon length (**d**) and colonic *Lcn2*, *Il1b*, *Il6* and *Tnf* mRNA levels (**e**) were analyzed on day 7. (**f**–**i**) B6.*Rag1*^−/−^ and NOD*.Rag1*^−/−^ mice were treated with 2.5% DSS for 7 days. Body weight changes (**f)** were monitored daily. Disease activity index (**g**), the colon length (**h**) and colonic *Lcn2*, *Il1b*, *Il6* and *Tnf* mRNA levels (**i**) were analyzed on day 7. Data represent the mean ± SEM. *n* = 6 mice per group. 2 independent experiments. \**p* < 0.05; \*\**p* < 0.01; \*\*\**p* < 0.001; \*\*\*\**p* < 0.0001; NS, not significant by log-rank test (**a**), two-way ANOVA with Sidak’s multiple comparison test (**b**, **f**) or 2-tailed Student’s *t* test (**c**–**e, g**–**i**).

Accumulating evidence indicates that the pathogenic loci commonly linked to T1D paradoxically protect against intestinal inflammation^18,19^. To unravel the immunological differences underlying the epidemiological separation of IBD and T1D, we modeled gut inflammation by treating B6 and NOD mice with DSS and evaluated their susceptibility to colitis. Indicators of colitis severity, such as body weight loss and the disease activity index (DAI) scores, were significantly lower in NOD mice than in B6 mice throughout the experimental course (Fig. 1b, c). This finding suggests that DSS administration produced a notably attenuated colitis phenotype in NOD mice compared with their B6 counterparts. Morphometric assessment showed relatively preserved colon length in NOD mice compared with B6 mice (Fig. 1d). Transcriptional profiling showed that the expression levels of key proinflammatory mediators, including lipocalin-2 (*Lcn2*), interleukin-1 beta (*Il1b*), interleukin-6 (*Il6*), and tumor necrosis factor-alpha (*Tnf*), in colonic tissues were significantly lower in NOD mice than in B6 mice after DSS administration (Fig. 1e). These results suggest that T1D-prone mice exhibited resilience to chemical-induced colitis, as evidenced by reduced clinical, histopathological, and molecular signs of inflammation.

To verify whether the adaptive immunity is critical for modulation of the protective phenotype of NOD mice to DSS-induced colitis, we examined the susceptibility of adaptive immunity-deficient B6.*Rag1*^−/−^ and NOD.*Rag1*^−/−^ mice to DSS-induced colitis. Disease indicators, including body weight loss, DAI, and colon lengths, were indistinguishable between B6.*Rag1*^−/−^ and NOD.*Rag1*^−/−^ mice throughout the experimental course (Fig. 1f–h). These results suggest that adaptive immunity is essential to the protective effect of DSS-induced colitis in NOD mice. The expression profile of colonic proinflammatory genes, including *Lcn2*, *Il1b*, *Il6*, and *Tnf*, showed no significant difference between B6.*Rag1*^−/−^ and NOD.*Rag1*^−/−^ mice after DSS administration (Fig. 1i), which supports the conclusion that IBD development is attenuated in NOD mice through mechanisms that operate in an adaptive immunity-dependent manner. Collectively, these findings suggest that the predisposition toward T1D-harboring intrinsic immunomodulatory properties in NOD mice confers a measurable reduction in susceptibility to DSS-induced colitis. This protective effect in NOD mice aligns with epidemiological evidence indicating lower incidence rates of IBD among individuals with T1D and suggests a pathophysiological link between clinical observations and experimental validation.

### IL-21 potentiates resistance of NOD mice to DSS-induced colitis

We next analyzed transcriptomic data of intestinal biopsies from patients with UC and non-IBD controls^32,33^. We found evidence of enrichment of *IL10* and *IL21* transcripts within the intestinal tissue of non-IBD controls (Fig. 2a). We also observed that the expression of *IL21* transcripts was significantly lower in the intestinal tissue of patients with UC than in non-IBD controls, whereas the expression levels of *IFNg*, *IL4*, and *IL10* transcripts were indistinguishable between these 2 groups (Fig. 2b). These findings reflect an inverse correlation between *IL21* expression and UC development in humans.

**Fig. 2.**
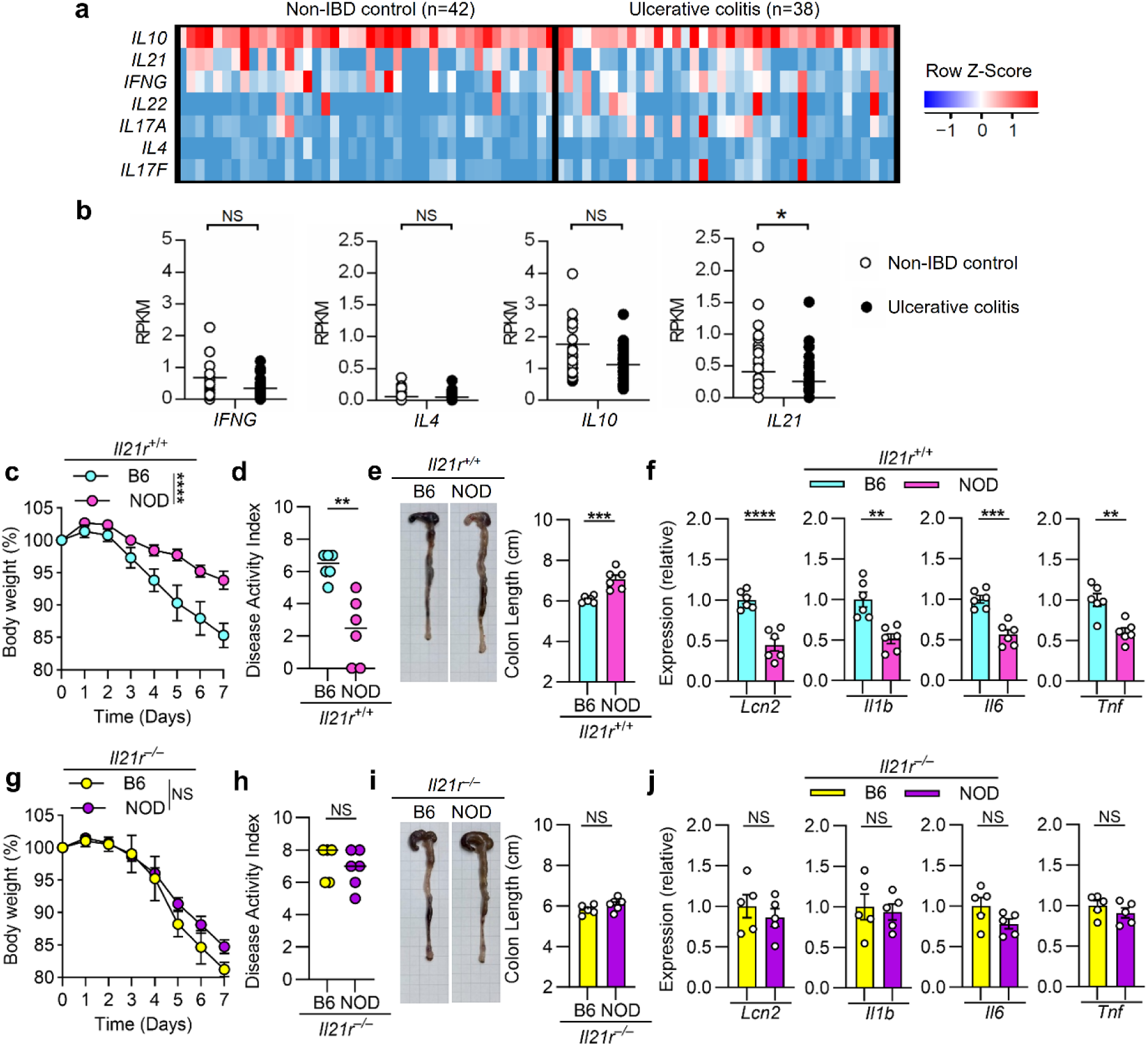
IL-21 potentiates resistance of NOD mice to DSS-induced colitis. (**a**) Gene expression profiling in human clinical samples. Heatmap displaying the expression of selected genes in ileal biopsies from non-IBD controls (n = 42) and patients with ulcerative colitis (n = 38). Data were retrieved from the GEO dataset GSE57945. (**b**) RPKM of *IFNg*, *IL4*, *IL10* and *IL21* in **a**. (**c**–**f**) B6.*Il21r*^+/+^ and NOD.*Il21r*^+/+^ mice were treated with 2.5% DSS for 7 days. Body weight changes (**c**) were monitored daily. Disease activity index (**d**), the colon length (**e**) and colonic *Lcn2*, *Il1b*, *Il6* and *Tnf* mRNA levels (**f**) were analyzed on day 7. (**g**–**j**) B6.*Il21r*^−/−^ and NOD.*Il21r*^−/−^ mice were treated with 2.5% DSS for 7 days. Body weight changes (**g**) were monitored daily. Disease activity index (**h**), the colon length (**i**) and colonic *Lcn2*, *Il1b*, *Il6* and *Tnf* mRNA levels (**j**) were analyzed on day 7. Data represent the mean ± SEM. *n* = 5-6 mice per group. 2 independent experiments. \**p* < 0.05; \*\**p* < 0.01; \*\*\**p* < 0.001; \*\*\*\**p* < 0.0001; NS, not significant by 2-tailed Student’s *t* test (**b**, **d**–**f**, **h**–**j**) or two-way ANOVA with Sidak’s multiple comparison test (**c**, **g**).

To investigate further whether IL-21 signaling is critical for the protection of NOD mice, we compared the susceptibility to DSS-induced colitis between B6.*Il21r*^−/−^ and NOD.*Il21r*^−/−^mice. Consistent with results described above (Fig. 1b–d), the disease indicators were significantly lower in NOD.*Il21r*^+/+^ mice than in B6.*Il21r*^+/+^ mice (Fig. 2c–e). Strikingly, these indicators were indistinguishable between B6.*Il21r*^−/−^ and NOD.*Il21r*^−/−^ mice throughout the experimental course (Fig. 2g–i). Taken together, these findings suggest a critical role of IL-21 in the protective phenotype in NOD mice. Colonic proinflammatory gene expression showed no significant differences between B6.*Il21r*^−/−^ and NOD.*Il21r*^−/−^ after DSS administration, whereas expression levels of these genes were significantly lower in NOD.*Il21r*^+/+^ mice than in B6.*Il21r*^+/+^ mice (Fig. 2f, j, respectively). These results confirm that NOD mice are protected from IBD induction through mechanisms that operate in an IL-21-signaling-dependent manner.

### An age-related reduction in c-Maf SUMOylation is associated with elevated IL-21 expression in CD4⁺ T cells from diabetes-susceptible mice

Transactivation of IL-21 in CD4⁺ T cells is regulated primarily by c-Maf ^34, 35^. Our previous report demonstrated that an age-associated reduction in c-Maf SUMOylation, rather than its expression level, critically increases *Il21* transactivation and contributes to autoimmune diabetes in NOD mice^36^. To explore further whether an age-related change in c-Maf SUMOylation also occurs and influences IL-21 expression in CD4⁺ T cells of nondiabetes-prone mice, we immunoprecipitated cell lysates of activated CD4^+^ T cells from B6 mice of different ages with anti-c-Maf followed by anti-SUMO-1 blotting. The ratio of SUMOylated to unSUMOylated c-Maf was significantly lower in CD4^+^ T cells from 12–14-week-old NOD mice than in CD4^+^ T cells from 12–14-week-old B6 mice. By contrast, the SUMOylated/unSUMOylated ratio was similar in CD4^+^ T cells from 6–8-week-old B6 and NOD mice (Fig. 3a). These findings suggest that an age-related reduction of c-Maf SUMOylation occurs in the NOD background.

**Fig. 3.**
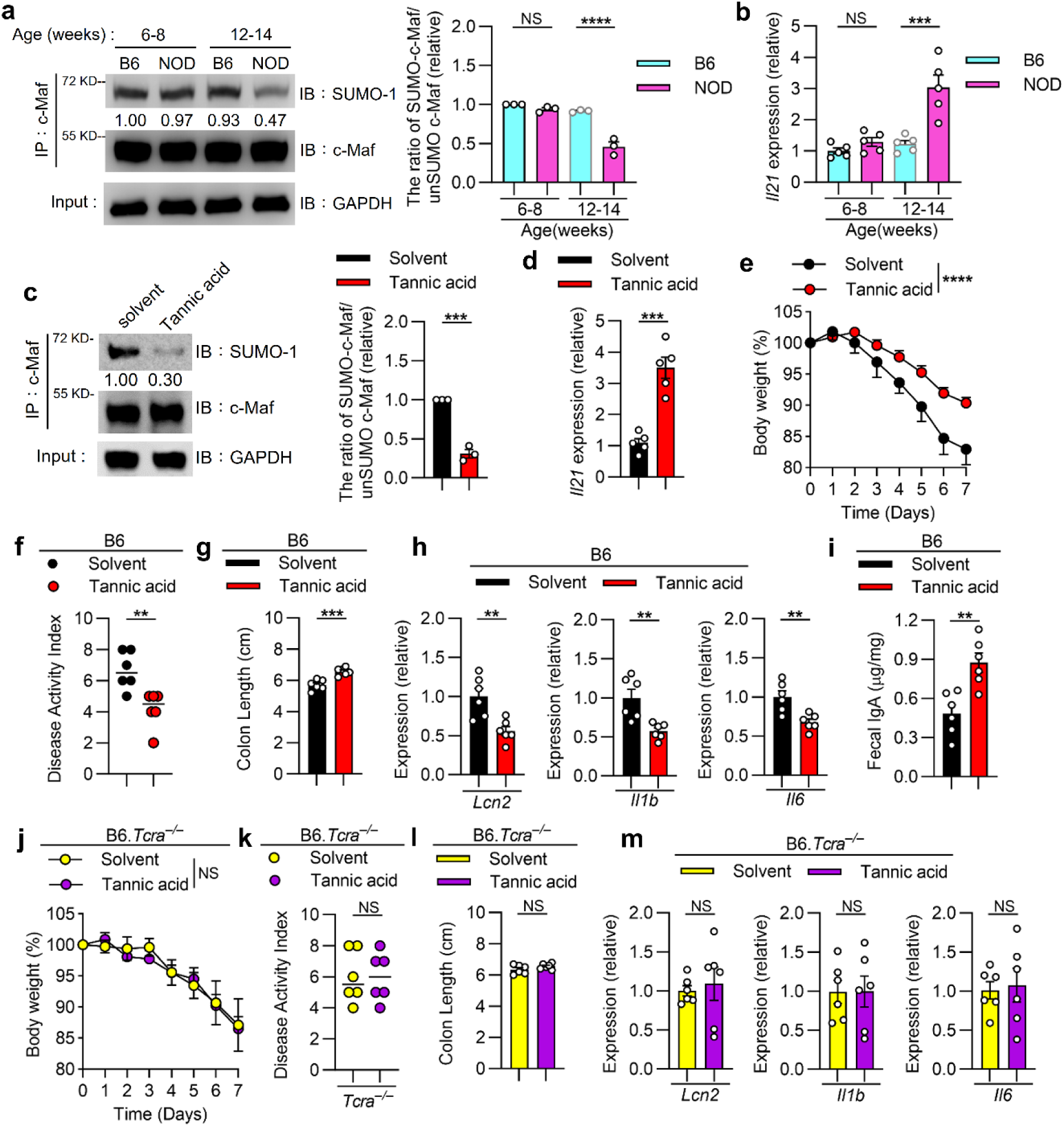
Impairment of c-Maf SUMOylation in CD4^+^ T cells induces the IL-21–IgA axis and ameliorates DSS-induced colitis. (**a**) Immunoprecipitation analysis of c-Maf SUMOylation in 6–8- and 12–14-week-old B6 and NOD CD4^+^ T cells cultured for 36 h with anti-CD3 and anti-CD28. (**b**) Expressions of *Il21* mRNA in CD4^+^ T cells cultured for 36 h as described in **a**. (**c**) Immunoprecipitation analysis of c-Maf SUMOylation in 6-8-week-old NOD CD4^+^ T cells cultured for 36 h with anti-CD3 and anti-CD28 in the presence of tannic acid (10 μM) or its solvent (PBS). (**d**) Expressions of *Il21* mRNA in CD4^+^ T cells as described in **c**. (**e**–**m**) B6.WT mice (**e**–**i**) or B6.*Tcra*^−/−^ mice (**j**–**m**) were treated with 2.5% DSS for 7 days and injected with solvent or tannic acid (20mg/Kg) daily. Body weight changes (**e**, **j**) were monitored daily. Disease activity index (**f**, **k**), the colon length (**g**, **l**), colonic mRNA levels (**h**, **m**) and fecal IgA levels (**i**) were analyzed on day 7. Data represent the mean ± SEM. *n* = 4-6 mice per group. 2 independent experiments. \*\**p* < 0.01; \*\*\**p* < 0.001; \*\*\*\**p* < 0.0001; NS, not significant by one-way ANOVA with Tukey’s multiple comparison test (**a**, **b**), 2-tailed Student’s *t* test (**c**, **d**, **g**–**i** and **k**–**m**) or two-way ANOVA with Sidak’s multiple comparison test (**e**, **j**).

We next examined the expression of IL-21 in CD4^+^ T cells from B6 and NOD mice at different ages. The expression of IL-21 mRNA was significantly higher in CD4^+^ T cells from 12–14-week-old NOD mice than in CD4^+^ T cells from 12–14-week-old B6 mice. By contrast, the expression of IL-21 mRNA was indistinguishable in CD4^+^ T cells from 6–8-week-old B6 and NOD mice (Fig. 3b). These data suggest an age-dependent inverse correlation between c-Maf SUMOylation and IL-21 production in CD4^+^ T cells from diabetes-prone mice.

### Tannic acid reduces c-Maf SUMOylation in CD4^+^ T cells and ameliorates DSS-induced colitis by promoting the IL-21–IgA axis

To investigate further whether pharmacological intervention in the SUMOylation process in CD4^+^ T cells from nondiabetic-prone mice affects c-Maf-modulated IL-21 production, we treated CD4^+^ T cells from B6 mice with tannic acid, which is a natural and nontoxic SUMOylation inhibitor ^37^. The amount of SUMOylated c-Maf was significantly lower in tannic acid-treated B6 CD4^+^ T cells than in solvent-treated cells, whereas the amount of unSUMOylated c-Maf was similar in both groups (Fig. 3c). We also found that the amount of IL-21 mRNA was significantly higher in tannic acid-treated B6 CD4^+^ T cells than in solvent-treated cells (Fig. 3d). These results indicate clearly that the pharmacological downregulation of c-Maf SUMOylation promotes its transactivation of *Il21* in CD4^+^ T cells from nondiabetes-prone mice.

To evaluate further whether tannic acid protects mice from development of colitis, we administered tannic acid to B6 mice and evaluated their susceptibility to DSS-induced colitis. The indicators of colitis severity and colonic proinflammatory gene expression were significantly lower in tannic acid-treated B6 mice than in solvent-treated mice throughout the experimental course (Fig. 3e, h). This finding suggests that the tannic acid-mediated reduction of c-Maf SUMOylation markedly attenuated the susceptibility to colitis in B6 mice.

Previous studies have shown that IL-21 is critical for mucosal immunity by promoting IgA class switching, which strengthens the intestinal barrier against inflammation, and that a deficiency in the IL-21 receptor (IL-21R) leads to impaired IgA responses and increased susceptibility to colitis ^38, 39, 40^. We found that fecal IgA level was significantly higher in tannic acid-treated B6 mice than in solvent-treated B6 mice (Fig. 3i), which supports a protective role of the IL-21–IgA axis in preventing DSS-induced colitis. To verify whether CD4^+^ T cells are critical for the tannic acid-mediated protection during colitis development, we examined the susceptibility of B6.*Tcra*^−/−^ mice with CD4^+^ T-cell deficiency to DSS-induced colitis. The disease indicators and colonic proinflammatory gene expression were indistinguishable between tannic acid-treated and solvent-treated B6.*Tcra*^−/−^ mice throughout the experimental course (Fig. 3j, m). These results suggest that CD4^+^ T cells are required for the tannic acid-mediated protection against DSS-induced colitis. Collectively, our findings demonstrate that SUMOylation-based intervention initiates the c-Maf-IL-21 axis in CD4⁺ T cells and triggers an IgA-based attenuation of the susceptibility to DSS-induced colitis.

### A T-cell-specific defect in c-Maf SUMOylation attenuates DSS-induced colitis via stimulating the IL-21–IgA axis

We next investigated whether a T-cell-restricted and c-Maf-based SUMOylation process modulates IL-21 production and suppresses colitogenesis in the DSS-induced mouse model. We first ablated endogenous c-Maf in T cells by establishing B6.*Maf^fl/fl^Lck^Cre^*mice, in which *loxP*-flanked *Maf* alleles (*Maf^fl/fl^*) are deleted by Cre recombinase driven by a T-cell-specific *Lck* promoter (*Lck^Cre^*) ^41^. These mice were then crossed with *Lck*-driven wild-type c-Maf (B6.*Lck^Maf-WT^*) or K33R c-Maf transgenic lines with a mutation at the SUMOylation site (B6.*Lck^Maf-K33R^*) to generate T-cell-specific B6.*Maf^WT^* and B6.*Maf^KR^* mice, respectively ^24^. We then treated these mice with DSS and compared the disease kinetics and severity. The indicators of colitis severity and colonic proinflammatory gene expression after DSS treatment were significantly lower in B6.*Maf^KR^*mice than in B6.*Maf^WT^* mice (Fig. 4a–d).

**Fig 4.**
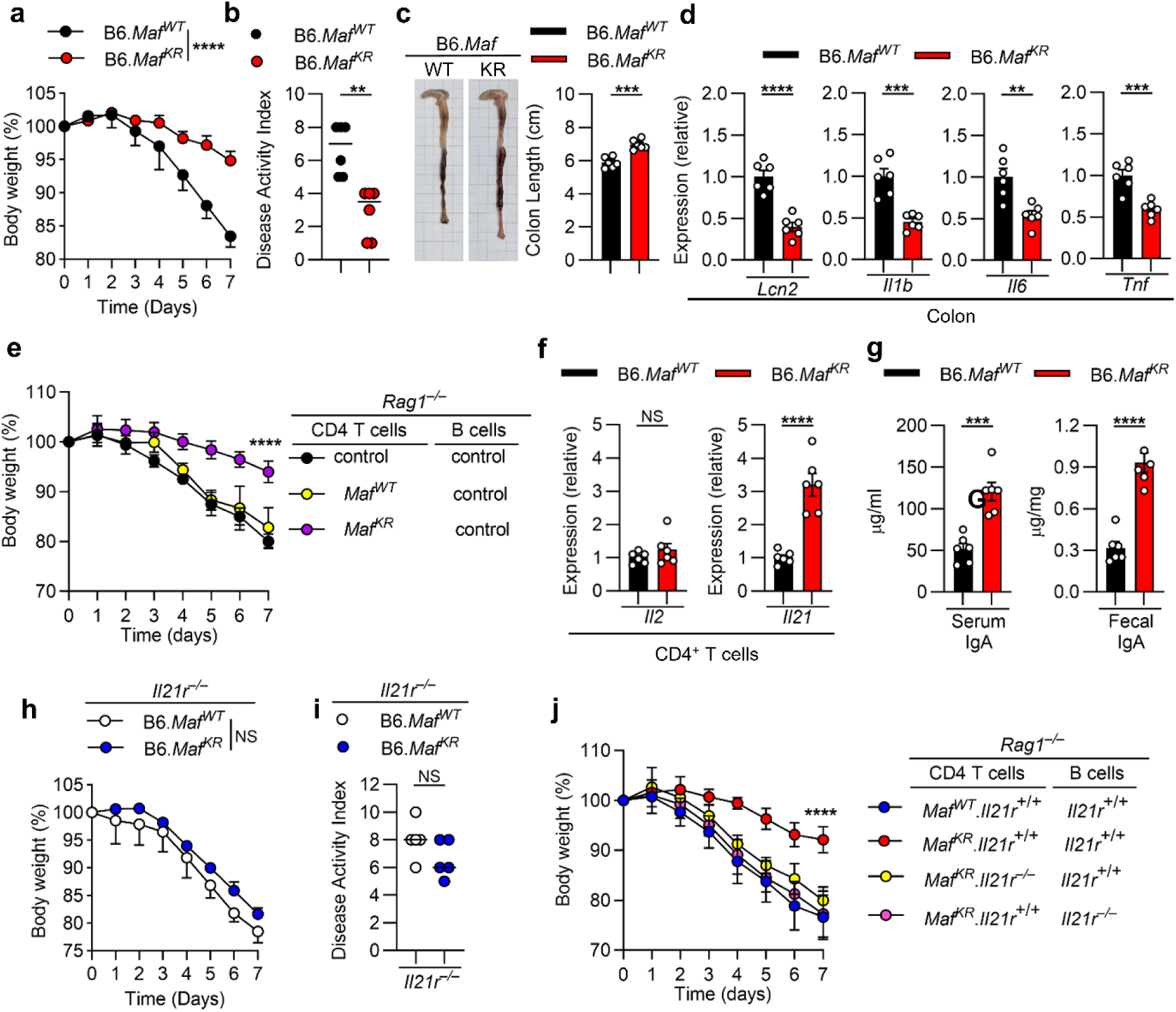
T-cell-specific c-Maf SUMOylation deficiency attenuates the severity of DSS-induced colitis via stimulating the IL-21–IgA axis. (**a**) B6.*MafWT* (B6.*Maffl/flLckCreLckMaf-WT*) and B6.*MafKR* (B6.*Maffl/flLckCreLckMaf-KR*) mice were treated with 2.5% DSS for 7 days. Body weight changes were monitored daily. (**b**–**d**) Disease activity index (**b**), the colon length (**c**) and colonic *Lcn2*, *Il1b*, *Il6* and *Tnf* mRNA levels (**d**) were analyzed on day 7 as in **a**. (**e**) B6.*Rag1*−/− mice were transferred with CD4+ T cells from indicated mice plus B220+ B cells from control mice (B6.*Maffl/fl*) for two weeks and were treated with 2.5% DSS for 7 days. Body weight changes were monitored daily. (**f**) Expressions of *Il2* and *Il21* mRNA in splenic CD4+ T cells were analyzed on day 7 as in **a**. (**g**) serum and fecal IgA levels were analyzed on day 7 as in **a**. (**h**) B6.*MafWTIl21r*–/– and B6.*MafKRIl21r*–/– mice were treated with 2.5% DSS for 7 days. Body weight changes were monitored daily. (**i**) Disease activity index were analyzed on day 7 as in **h**. (**j**) B6.*Rag1*−/− mice were transferred with CD4+ T cells plus B220+ B cells from indicated mice for two weeks and were treated with 2.5% DSS for 7 days. Body weight changes during DSS treatment were monitored daily. Data represent the mean ± SEM. *n* = 5-6 mice per group. 2 independent experiments. \*\**p* < 0.01; \*\*\**p* < 0.001; \*\*\*\**p* < 0.0001; NS, not significant by two-way ANOVA with Sidak’s multiple comparison test (**a**, **e**, **h**, **j**) or 2-tailed Student’s *t* test (**b**–**d**, **f**, **g**, **i**).

Given that *Lck*-driven Cre recombinase and Maf transgene expression affect both CD4^+^ and CD8^+^ T cells, we next examined whether CD4^+^ T cells are mainly responsible for this *Maf^KR^*-modulated protection against colitis. We cotransferred CD4^+^ T cells from the relevant donors plus control B cells into B6.*Rag1*^−/−^ recipients and monitored their disease kinetics after DSS administration. The mice that received *Maf^KR^* CD4^+^ T cells plus control B cells developed colitis more slowly than those that received control or *Maf^WT^* CD4^+^ cells plus control B cells (Fig. 4e), which suggests that the SUMO-defective c-Maf-mediated protection in *Maf^KR^*mice is CD4^+^ T-cell autonomous. We also analyzed CD4^+^ T cells from these 2 transgenic lines and found that the levels of *Il21* transcripts were markedly higher in B6.*Maf^KR^* CD4^+^ T cells than in B6.*Maf^WT^* CD4^+^ T cells but that their *Il2* transcript levels were indistinguishable (Fig. 4f). Strikingly, the levels of serum and fecal IgA were significantly higher in B6.*Maf^KR^* mice than in B6.*Maf^WT^* mice (Fig. 4g). Collectively, our results suggest that intervention in SUMOylation in a T-cell-specific and c-Maf-based manner triggers the IL-21–IgA axis, which attenuates the development of DSS-induced colitis.

To investigate whether the T-cell-specific/SUMO-defective c-Maf-mediated protection against DSS-induced colitis is IL-21R-signaling dependent, we crossed B6.*Maf^WT^* and B6.*Maf^KR^* mice with B6.*Il21r*^−/−^mice to generate B6.*Maf^WT^Il21r*^−/−^ and B6.*Maf^KR^Il21r*^−/−^ mice, respectively, and we monitored colitogenesis after DSS administration. In the absence of IL-21 signaling, B6.*Maf^KR^* mice lost their protection against colitis induction (Fig. 4h, i), which suggests an essential role of IL-21 in *Maf^KR^*-mediated protection. To dissect the necessity of IL-21R signaling for disease protection induced by CD4^+^ T and/or B cells, we cotransferred CD4^+^ T cells from B6.*Maf^KR^*.*Il21r*^+/+^ or B6.*Maf^KR^*.*Il21r*^−/−^ mice with B cells from B6.*Il21r*^+/+^ or B6.*Il21r*^−/−^ mice into B6.*Rag1*^−/−^ mice and monitored their colitis kinetics after DSS administration. The body weight loss was greater in mice that received *Maf^KR^*.*Il21r*^−/−^ CD4^+^ plus *Il21r*^+/+^ B cells or *Maf^KR^*.*Il21r*^+/+^ CD4^+^ plus *Il21r*^−/−^ B cells than in mice that received *Maf^KR^*.*Il21r*^+/+^ CD4^+^ plus *Il21r*^+/+^ B cells (Fig. 4j). This finding suggests that IL-21 signaling is required by both CD4^+^ T cells and B cells to mediate protection against the development of colitis.

### SUMO-defective c-Maf in T cells reshapes the gut microbiota and metabolic profile toward an LCA-rich environment

Previous studies have reported that IgA maintains intestinal homeostasis by physically enchaining proliferating bacteria and functionally shaping the composition and metabolic activity of the gut microbiota to promote symbiosis ^42, 43^. To investigate whether the SUMO-defective c-Maf-triggered IL-21–IgA axis attenuates the development of DSS-induced colitis by modulating the intestinal microbiota, we conducted full-length 16S ribosomal RNA gene sequencing of fecal samples collected from B6.*Maf^WT^* and B6.*Maf^KR^* mice and comprehensively characterized and compared the gut microbial communities. The top 10 bacterial species were quantified based on their relative abundance in fecal samples using comparative analysis of gut bacterial abundance of B6.*Maf^WT^* and B6.*Maf^KR^* mice (Fig. 5a). The abundance of *Lactobacillus johnsonii* (*L. johnsonii*) was higher but that of *Dubosiella newyorkensis* (*D. newyorkensis*) was markedly lower in B6.*Maf^KR^* mice than in B6.*Maf^WT^* mice (Fig. 5b). The abundance of the other species, including *Muribaculum intestinale* (*M. intestinale*), *Bacteroides intestinihominis* (*B. intestinihominis*), *Flavonifractor rodentium* (*F. rodentium*), *Klebsiella alsydes* (*K. alsydes*), *Clostridium disporicum* (*C. disporicum*), *Roseburia champanellensis* (*R. champanellensis*), *Enterocloster massiliensis* (*E. massiliensis*) and *L. reuteri* did not differ significantly between the 2 groups. These findings suggest that the SUMO-defective c-Maf-triggered IL-21–IgA axis modulates specific microbial taxa and might thereby contribute to altered intestinal immune or metabolic states.

**Fig 5.**
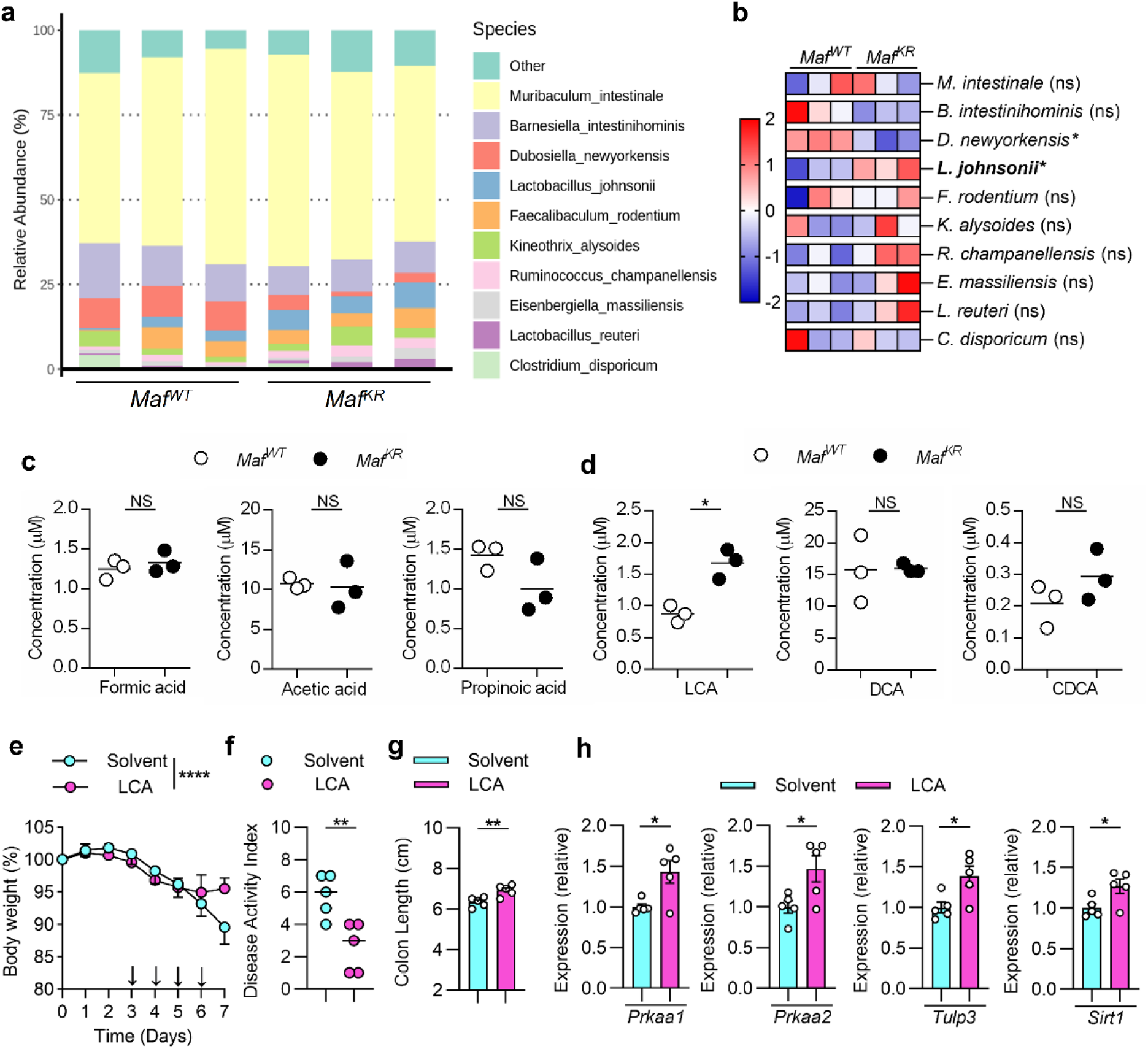
A T-cell-specific defect in c-Maf SUMOylation remodels the gut microbiota to promote *Lactobacillus johnsonii* expansion and LCA production. (**a**) Fecal DNA from B6.*Maf^WT^* and B6.*Maf^KR^* mice was analyzed by full-length 16S rRNA sequencing to identify the top ten most abundant species. (**b**) Relative abundance of the top 10 bacterial species in fecal microbiota from B6.*Maf^WT^* and B6.*Maf^KR^* mice as in **a**. (**c**, **d**) Targeted metabolomic analysis of fecal samples from B6.*Maf^WT^* and B6.*Maf^KR^* mice by mass spectrometry. Panels show the concentrations of (**c**) short-chain fatty acids (formic, acetic, and propionic acid) and (**d**) bile acids (LCA, DCA, and CDCA). (**e**–**h**) B6 mice were treated with 2.5% DSS for 7 days and injected with solvent or LCA (10mg/Kg) daily from day 3 to day 6. Body weight changes (**e**) were monitored daily. Disease activity index (**f**), the colon length (**g**), colonic *Prkaa1 Prkaa2*, *Tulp3* and *Sirt1* mRNA levels (**h**) were analyzed on day 7. Data represent the mean ± SEM. *n* = 3-5 mice per group. 2 independent experiments. \**p* < 0.05; \*\**p* < 0.01; \*\*\*\**p* < 0.0001; NS, not significant by 2-tailed Student’s *t* test (**b**–**d** and **f**–**g**) or two-way ANOVA with Sidak’s multiple comparison test (**e**).

We next performed quantitative analysis of fecal metabolites to compare the concentrations of 6 representative acids such as short-chain fatty acids (SCFAs) and bile acids (BAs) between the 2 groups. The concentrations of SCFAs, including formic acid, acetic acid, and propionic acid, were similar in the 2 groups (Fig. 5c). By contrast, BA profiling revealed significantly higher concentration of LCA in B6.*Maf^KR^* mice than in B6.*Maf^WT^* mice, whereas the levels of deoxycholic acid (DCA) and chenodeoxycholic acid (CDCA) were similar in the 2 groups. (Fig. 5d). These results suggest that the SUMO-defective c-Maf-triggered IL-21–IgA axis promotes shifts in microbial BA metabolism and thereby contributes to the protection against DSS-induced colitis.

Recent findings underscore that the metabolic benefits of LCA are regulated by the AMPK signaling axis^44,45^. To determine whether this specific pathway is involved in the efficacy of LCA in preventing or treating colitis, we administered LCA to B6 mice and evaluated the disease severity of DSS-induced colitis. The indicators of colitis severity were significantly lower in LCA-treated B6 mice than in solvent-treated mice throughout the experimental course (Fig. 5e–g), which suggests that LCA treatment ameliorated the pathogenesis of DSS-induced colitis. LCA also significantly increased the intestinal mRNA expression of the core components in the AMPK-signaling axis including *Prkaa1*, *Prkaa2*, *Tulp3*, and *Sirt1* (Fig. 5h). These findings support the idea that LCA suppresses DSS-induced colitis by activating this specific regulatory pathway, which subsequently induces the anti-inflammatory effects mediated by the AMPK-signaling cascade.

### The HDAC2 inhibitor BRD6688 promotes c-Maf-mediated IL-21 expression and attenuates intestinal inflammation

Our previous study reported that the SUMOylation status of c-Maf critically dictates the epigenetic regulation of *Il21* transcription by inducing the recruitment of HDAC2 to the promoter region^24^. To evaluate whether pharmacological targeting of HDAC2 provides protection through the c-Maf–IL-21–IgA axis in DSS-induced colitis, we administered the HDAC2 inhibitor BRD6688 to B6 mice^46^ and evaluated the severity of DSS-induced colitis. The indicators of colitis severity and colonic proinflammatory gene expression were significantly lower in BRD6688-treated B6 mice than in solvent-treated mice throughout the experimental course (Fig. 6a–d), which suggests that BRD6688-mediated HDAC2 modulation plays a protective role against colitis induction.

**Fig 6.**
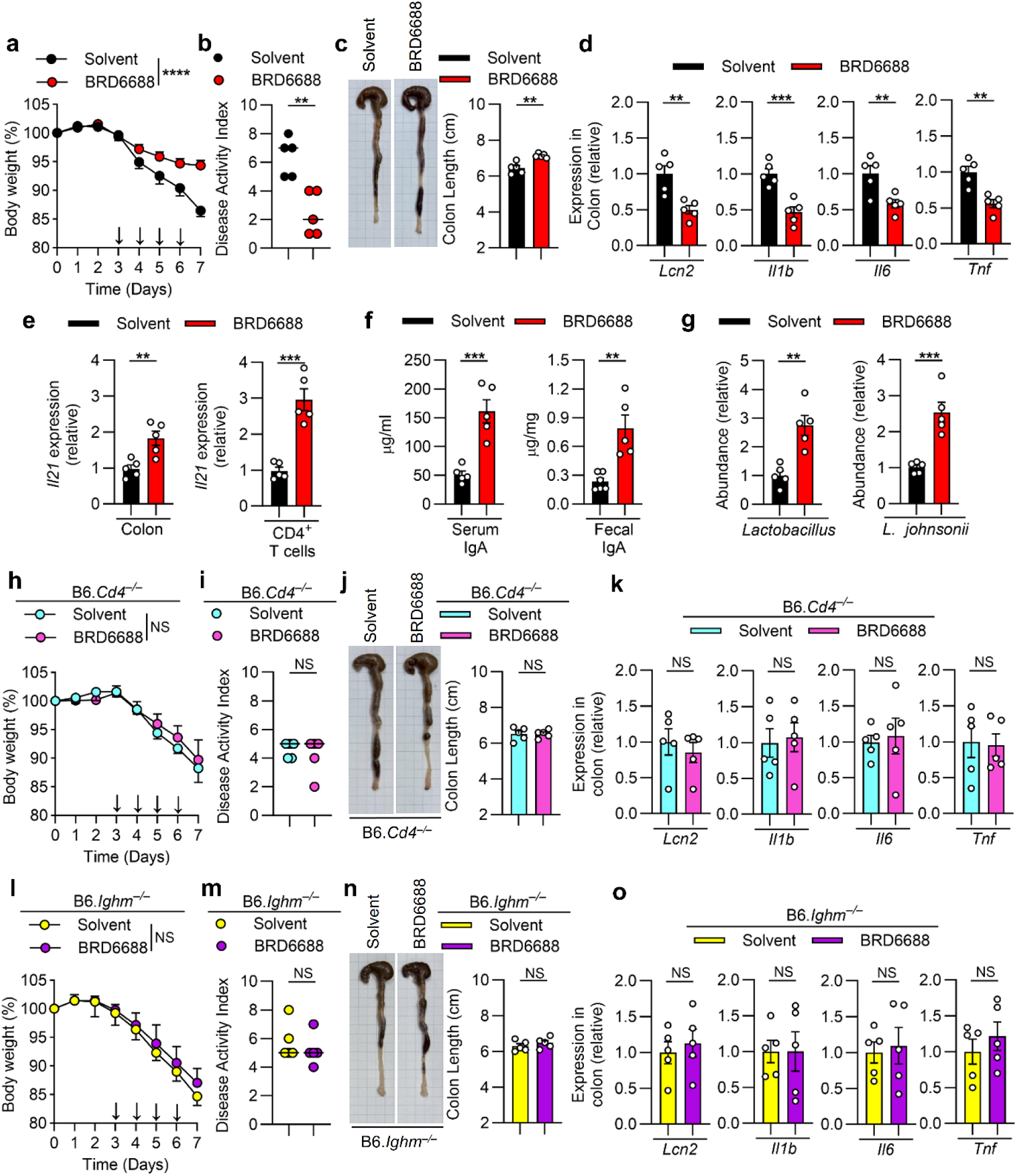
The HDAC2 inhibitor BRD6688 promotes c-Maf-mediated IL-21 expression and attenuates intestinal inflammation. (**a**–**g**) B6.WT, (**h**–**k**) B6.*Cd4*^−/−^ mice and (**l**–**o**) B6.*Ighm*^−/−^ mice were treated with 2.5% DSS for 7 days and injected with solvent or BRD6688 (5mg/Kg) daily from day 3 to day 6. Body weight changes (**a**, **h**, **l**) were monitored daily. Disease activity index (**b**, **i**, **m**), the colon length (**c**, **j**, **n**), colonic mRNA levels (**d**, **k**, **o**), *Il21* mRNA levels in colon tissues and CD4^+^ T cells (**e**), serum and fecal IgA levels (**f**) and relative abundance of *Lactobacillus* and *L. johnsonii* in fecal DNA (**g**) were analyzed on day 7. Data represent the mean ± SEM. *n* = 5-6 mice per group. 2 independent experiments. \*\**p* < 0.01; \*\*\**p* < 0.001; \*\*\*\**p* < 0.0001; NS, not significant by two-way ANOVA with Sidak’s multiple comparison test (**a**, **h**, **l**) or 2-tailed Student’s *t* test (**b**–**g**, **i**–**k. m**–**o**).

We next analyzed colon tissues and CD4^+^ T cells from these 2 groups of mice and found that the Il21 transcript level was markedly higher in the BRD6688-treated group than in the solvent-treated group (Fig. 6e). Serum and fecal IgA levels were significantly higher and *L. johnsonii* was significantly enriched (Fig. 6f, g) in BRD6688-treated mice. These findings suggest that BRD6688 promotes both mucosal immune responses and beneficial microbial shifts.

To delineate the cellular compartments responsible for the therapeutic effects of BRD6688, we investigated whether the adaptive immune system, specifically CD4^+^ T cells and B cells, is required for the protection observed in the DSS-induced colitis model. The protective benefits of BRD6688 observed in wild-type mice were abrogated in B6.*Cd4*^−/−^ mice. Throughout the DSS challenge, BRD6688-treated B6.*Cd4*^−/−^ mice exhibited severe clinical symptoms, including drastic weight loss and elevated DAI scores, which mirrored the phenotypes of their solvent-treated counterparts (Fig. 6h–k). Parallel experiments using B-cell-deficient B6.*Ighm*^−/−^ mice yielded similar results in that BRD6688 failed to rescue the intestinal architecture or reduce inflammation in the absence of B cells (Fig. 6l–o). These genetic loss-of-function data demonstrate clearly that BRD6688 requires a synergistic CD4^+^ T-cell and B-cell-dependent mechanism to mitigate colonic inflammation. Collectively, our findings demonstrated that SUMOylation-based intervention in CD4⁺ T cells initiates the c-Maf-IL-21 axis and triggers an IgA-based attenuation of DSS-induced colitis.

### Pharmacological inhibition of HDAC2 suppresses PBMC-derived colitis

To assess further the translational potential of the manipulation of the c-Maf SUMOylation–HDAC2–IL-21 axis in colitis, we established a human PBMC-derived colitis model and evaluated the therapeutic efficacy of BRD6688. Triple-knockout (TKO) B6.*Rag2*^−/−^*Cd47*^−/−^*Il2rg*^−/−^ mice were reconstituted with human PBMCs obtained from 2 independent healthy donors (n = 3 mice per donor). Following a 3-week engraftment period to allow for human immune system reconstitution, colitis was induced by administering DSS for 7 days; the mice received daily intraperitoneal injections of either solvent or BRD6688 from day 3 to 6 (Fig. 7a). BRD6688 treatment substantially suppressed the production of key proinflammatory cytokines and significantly reduced inflammatory damage in mice reconstituted from both Donors #1 and #2 (Fig. 7b–i). This parallel improvement across different donor-derived cohorts demonstrates that the protective effects of BRD6688 are robust against interindividual variability. Collectively, our results demonstrate that pharmacological interference with HDAC2 builds on the protective mechanism, which underscores the strong translational potential of HDAC2 inhibition as a therapeutic strategy for IBD.

**Fig. 7.**
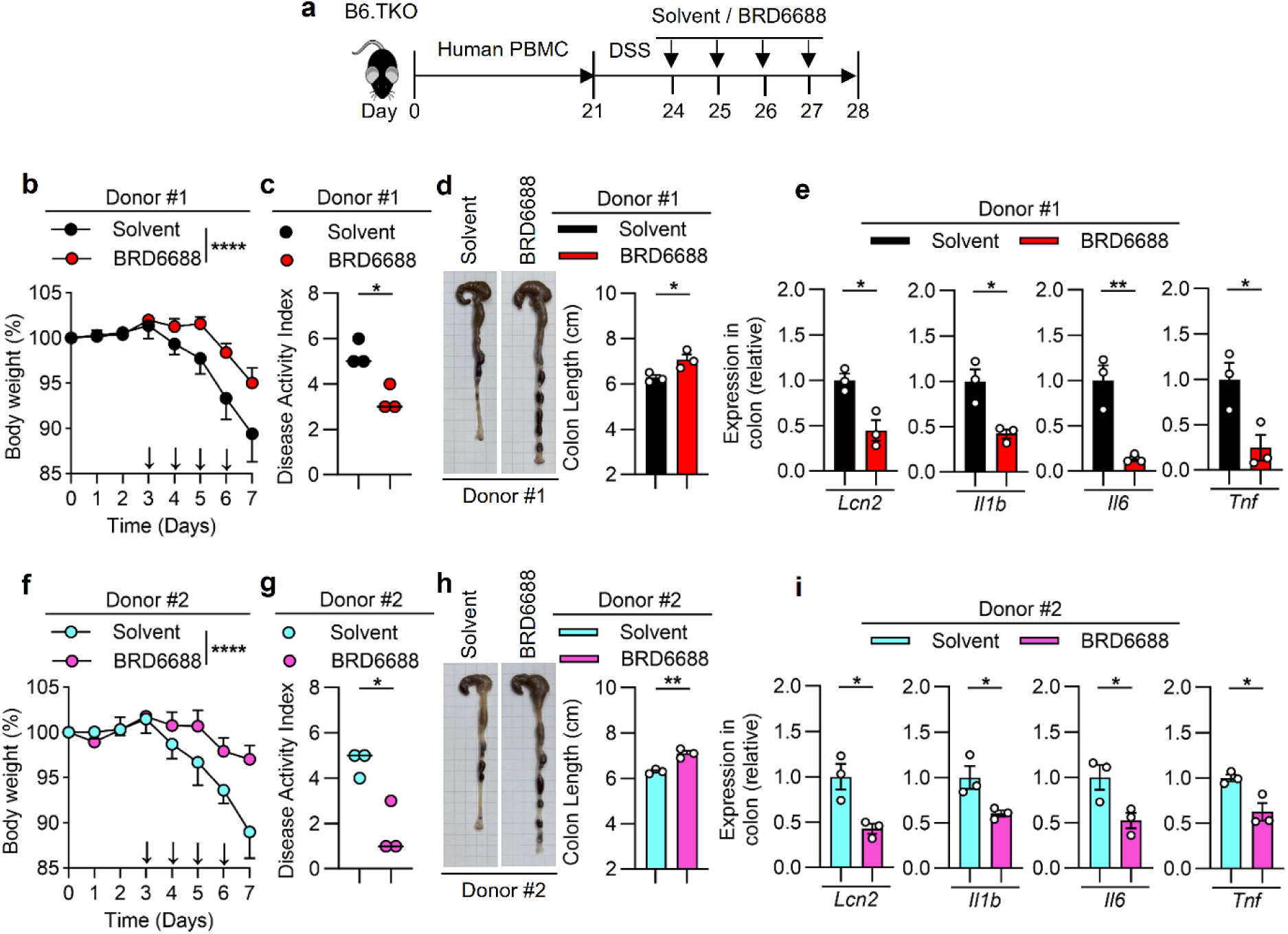
Therapeutic efficacy of the HDAC2 inhibitor BRD6688 in a PBMC-humanized mouse model of colitis. (**a**) B6.*Rag2*^−/−^*Cd47*^−/−^*Il2rg*^−/−^ (triple-knockout, TKO) mice were transplanted with human PBMCs derived from two independent donors: Donor #1 (**b**–**e**) and Donor #2 (**f**–**i**). After 21 days of engraftment, mice were treated with 2.5% DSS for 7 days and injected with solvent or BRD6688 (5mg/Kg) daily from day 24 to day 27. (**b**, **f**) Body weight changes were monitored daily. Disease activity index (**c** and **g**), the colon length (**d**, **h**), colonic *Lcn2*, *Il1b*, *Il6* and *Tnf* mRNA levels (**e**, **i**) were analyzed on day 7. Data represent the mean ± SEM. *n* = 3 mice per group. 2 independent experiments. \**p* < 0.05; \*\**p* < 0.01; \*\*\*\**p* < 0.0001; NS, not significant by two-way ANOVA with Sidak’s multiple comparison test (**b**, **f**) or 2-tailed Student’s *t* test (**c**–**e** and **g**–**i**).

In conclusion, our study identifies a previously unrecognized immunological crosstalk in which a single transcription factor-based modification dictates the susceptibility to both T1D and IBD. By integrating a 14-year population-based cohort study with detailed mechanistic inquiries in T-cell-specific c-Maf SUMOylation-defective mice, we have elucidated a robust T cell–microbiota–metabolite axis. As synthesized in our working model (Fig. 8), the impaired c-Maf SUMOylation is driven by its interaction with HDAC2, which licenses an enhanced IL-21–IgA regulatory program. This transcriptional shift promotes the enrichment of *L. johnsonii* and subsequent elevation of the concentration of the microbial metabolite LCA, which collectively trigger the AMPK anti-inflammatory pathway within the intestinal microenvironment. The successful application of the HDAC2 inhibitor BRD6688 in our human PBMC-derived colitis model demonstrates that this endogenous protective circuit can be pharmacologically harnessed. These findings provide a mechanistic explanation for the epidemiological resistance to IBD in T1D patients, highlights the c-Maf SUMOylation–HDAC2 axis as a high-value therapeutic target, and paves the way for a new generation of precision immunotherapies for IBD.

**Fig. 8.**
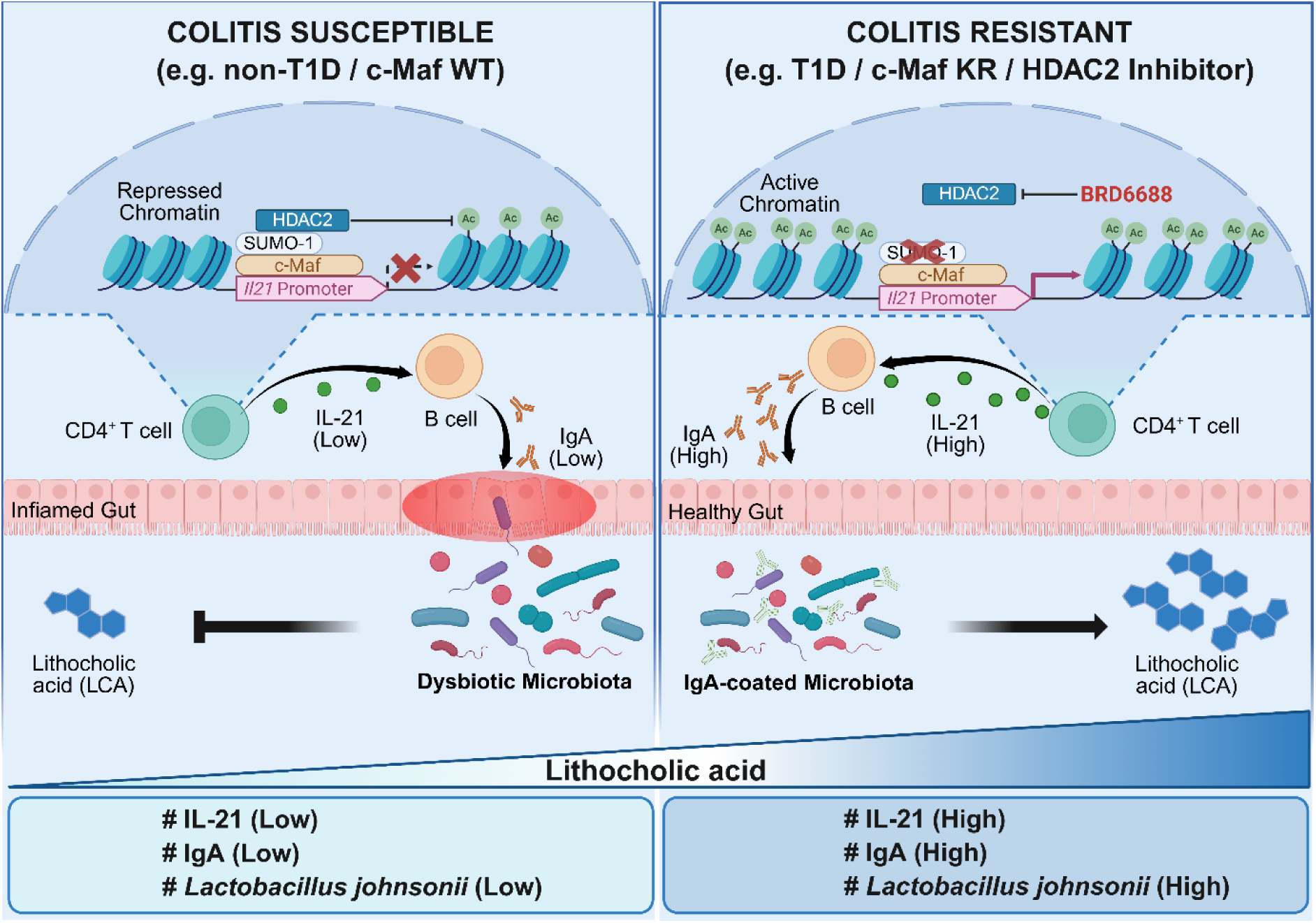
Proposed mechanistic model of the c-Maf SUMOylation molecular switch in modulating IBD susceptibility. (Left) Colitis susceptible state: In the absence of T1D or in c-Maf WT conditions, HDAC2-mediated deacetylation and c-Maf SUMOylation lead to repressed chromatin at the *Il21* promoter, resulting in low IL-21 and IgA levels. This fosters a dysbiotic microbiota and low LCA levels, predisposing the host to gut inflammation. (**Right**) Colitis resistant state: In T1D-associated conditions or via pharmacological HDAC2 inhibition (BRD6688), c-Maf SUMOylation is abrogated, and chromatin remains active. This “molecular switch” triggers high IL-21 and IgA production, leading to IgA-coated microbiota, increased *L. johnsonii* abundance, and robust LCA production, which maintains a healthy gut environment and confers resistance to colitis.

## Discussion

Our study reveals a distinct inverse comorbidity between T1D and IBD that challenges the conventional paradigm that autoimmune susceptibilities universally coaggregate. Leveraging a comprehensive 14-year population-based cohort of over 139,000 patients, we have shown that the organ-specific autoimmune background of T1D paradoxically confers significant resistance to intestinal inflammation. This finding aligns with recent Mendelian randomization studies^19^ and suggests a potential protective causal effect of T1D genetic liability on the risk of UC. We have identified the molecular basis of this inverse comorbidity as a distinct posttranslational defect of a single transcription factor in T cells. Under physiological condition, c-Maf SUMOylation may function as a critical licensing step that enables the initiation of intestinal inflammation. In the context of the T1D in the NOD mouse model, the inherent deficiency of this modification serves as a protective brake on colitis development. This SUMOylation-based alteration reshapes the mucosal immune landscape and redirects it toward a state that resists inflammatory pathology.

Expanding on our previous finding that c-Maf SUMOylation recruits HDAC2 to repress *Il21* during diabetogenesis^24^, we have shown here that the genetic abrogation of this recruitment in T1D sustains a hyperacetylated and transcriptionally permissive *Il21* locus in colonic CD4^+^ T cells. This permissive state drives increased IL-21 production that is prerequisite for a robust protective IgA response. Thus, the c-Maf-HDAC2–IL-21–IgA axis constitutes a precise host-intrinsic rheostat through which the specific posttranslational modification dictates the threshold of colitis susceptibility. The broad impact of epigenetic remodeling by HDACs in regulating mucosal integrity is well recognized^47,48,49^, and we have identified c-Maf SUMOylation as the specific molecular address directing HDAC2 to the *Il21* promoter to control gut inflammation. Our study findings expand the biological paradigm of our previous findings by demonstrating that the host repurposes the same c-Maf and HDAC2 repression machinery used in organ-specific autoimmunity to license mucosal barrier defense and provides a unified mechanistic basis for the inverse comorbidity between T1D and IBD.

Beyond the host-intrinsic machinery, we have also demonstrated a crucial host–microbe metabolic handoff that seems to amplify this protection. We found that an IgA-rich environment fostered by c-Maf SUMOylation deficiency favors the expansion of the BA-metabolizing commensal *L. johnsonii*, which is consistent with the established role of IgA in facilitating the stable colonization of beneficial mucosal symbionts ^50^. Building on recent findings identifying LCA as a TULP3-dependent activator of the sirtuin–AMPK axis^44,45^, we propose a dual-hit model by which c-Maf SUMOylation deficiency-driven IgA remodels the microbiota to increase LCA production. This finding integrates our work with emerging concepts in immunometabolism by suggesting that LCA serves as a key messenger that bridges the gap between microbial metabolism and host tissue resilience. Such a view is consistent with a protective role of *L. johnsonii*-derived BAs reported in other inflammatory contexts^51^.

Translating these mechanistic insights into a therapeutic strategy, we provide proof-of-concept evidence that this protective phenotype can be pharmacologically recapitulated in nondiabetic hosts. By administering the HDAC2 inhibitor BRD6688, we successfully mimicked the transcriptional signature of c-Maf SUMOylation deficiency to restore *Il21* promoter acetylation and suppress colitis severity in vivo. Our results suggest that the c-Maf–HDAC2 axis is a viable and druggable target. Identification of the *L. johnsonii*–LCA–AMPK pathway opens avenues for combinatorial therapies. Given the potent anti-inflammatory effects of AMPK activation in the gut, future regimens might pair epigenetic modulators such as HDAC2 inhibitors with precision probiotics or LCA mimetics to maximize therapeutic efficacy in treating IBD.

Although our study establishes a robust mechanistic framework, we acknowledge that the epidemiological association in humans requires further validation through prospective studies that integrate patient metabolomics to confirm a clinical LCA–AMPK link. Additionally, the broad effects of HDAC inhibition necessitate the future development of highly selective agents or cell type-specific delivery systems to minimize off-target risks. In conclusion, we have reported that a single transcription factor-based SUMOylation deficiency in T cells can be repurposed by the host to license gut protection. By elucidating this reciprocal modulation mediated by c-Maf SUMOylation and the LCA–AMPK metabolic circuit, our study highlights a novel therapeutic potential for treating IBD.

## Methods

### Data source and study design

This nationwide, population-based retrospective cohort study used the National Health Insurance Research Database (NHIRD) in Taiwan, which provides a universal single-payer system covering >99% of the population^52,53^. The study protocol was approved by the Institutional Review Board of Chang Gung Memorial Hospital (201900566B0). Although the source data retrieved covered the period from January 1, 1997, to December 31, 2013, the initial 3-year period (1997–1999) was used strictly to screen for historical medical records and exclude preexisting conditions to ensure diagnostic stability. Consequently, the primary analytical timeframe for ascertaining incident cases spanned 14 years, from January 1, 2000, to December 31, 2013.

### Disease definitions and validation

To ensure rigorous diagnostic accuracy, patients with T1D and IBD were identified based on strict criteria: (1) at least 2 outpatient visits or 1 inpatient admission with disease-specific diagnoses (T1D: ICD-9-CM 250.X1/250.X3; CD: 555.XX; UC: 556.XX)^54,55^ and (2) possession of a catastrophic illness certificate (CIC). The CIC serves as a validation gold standard within the NHIRD because its application requires a comprehensive review of clinical data, pathology reports, and imaging by independent senior specialists (endocrinologists or gastroenterologists) commissioned by the NHI Administration. This strict verification process minimizes coding errors and ensures high data validity.

### Exclusion criteria and statistical matching

To evaluate specifically the risk of IBD development following T1D onset, we excluded patients diagnosed with IBD before their T1D diagnosis or the index date. Additional exclusion criteria included malignancy, AIDS, or incomplete demographic data. To minimize selection bias, we implemented a 1:1 propensity score matching strategy^56,57^ to align the T1D cohort with a nondiabetic control group based on age, sex, and multiple comorbidities. All participants were followed longitudinally from the index date until the diagnosis of IBD, withdrawal from the NHI program, or the end of the study (December 31, 2013).

### Mice

NOD/Sytwu (K^d^, D^b^, I-A^g7^, I-E^null^), NOD.*Rag1*^−/−^, B6.*Rag1*^−/−^, B6.*Tcra*^−/−^, B6.*Il21r*^−/−^, B6.*Cd4*^−/−^, B6.*Ighm*^−/−^ and B6.*Rag2*^−/−^*Cd47*^−/−^*Il2rg*^−/−^ mice were purchased from The Jackson Laboratory. Wild-type C57BL/6 mice were obtained from the National Institutes of Applied Research National Center for Biomodels (Taipei, Taiwan). B6.*Maf^fl/fl^* mice were kindly provided by Professor Shi-Chuen Miaw (National Taiwan University, Taipei, Taiwan). B6.*Lck^Cre^*, B6.*Lck^Maf-WT^*, B6.*Lck^Maf-K33R^*, and NOD.*Il21r*^−/−^mice were established as previously described^17,24,58^. All mice were housed and bred under specific-pathogen-free conditions at the animal center of the National Defense Medical University (Taipei, Taiwan).

### DSS-induced colitis model

Colitis was induced by administering 2% (w/v) DSS (36–50 kDa; MP Biomedicals, Irvine, CA, USA) in drinking water provided *ad libitum*. Disease severity was monitored daily using a modified clinical scoring system based on a previous report^59^. The DAI was calculated by summing the scores for weight loss (0, none; 1, 1–5%, 2, 5–10%; 3, 10–15%; 4,: >15%), stool consistency (0, normal; 2, loose stools; 4, watery diarrhea), and rectal bleeding (0, normal; 2, slight bleeding; 4, gross bleeding).

### Cell isolation and activation

Splenic CD4^+^ T cells and B220^+^ B cells were isolated from the indicated mouse strains using magnetic separation kits (BioLegend, San Diego, CA, USA) according to the manufacturer’s protocols. Post-sort purity was consistently >95% as determined by flow cytometry. For *in vitro* stimulation, cells were cultured in wells precoated with anti-CD3 antibody (clone 145-2C11; 5 μg/mL; BD Biosciences, Franklin Lakes, NJ, USA) in the presence of soluble anti-CD28 antibody (clone 37.51; 2 μg/mL; BioLegend).

### Chemicals and reagents

Tannic acid and *N*-ethylmaleimide (NEM) were purchased from Sigma-Aldrich (St. Louis, MO, USA). LCA and the HDAC2 inhibitor BRD6688 were obtained from MedChemExpress (Monmouth Junction, NJ, USA).

### Reverse transcription and quantitative real-time PCR

Total RNA was extracted using RNeasy Mini Kits (Qiagen, Hilden, Germany) and reverse transcribed into cDNA using the SuperScript III First-Strand Synthesis System (Invitrogen, Carlsbad, CA, USA) according to the manufacturer’s instructions. Quantitative real-time PCR was performed using SYBR Green PCR Master Mix on an QuantStudio 7 Flex Real-Time PCR System (Applied Biosystems, Waltham, MA, USA). Expression was normalized to the expression of the housekeeping gene *Rps29*.

### Immunoprecipitation and western blot analysis

CD4^+^ T cells were lysed using RIPA buffer (50 mM Tris, pH 7.4, 2% SDS, 150 mM NaCl) containing a protease inhibitor mixture (Sigma-Aldrich) on ice for 30 min. For detection of SUMO-conjugated c-Maf, CD4^+^ T cells were washed twice with ice-cold PBS supplemented with 20 mM NEM and lysed in SDS lysis buffer (5% SDS, 0.15 M Tris-HCl pH 6.7, and 30% glycerol) diluted 1:2 in RIPA buffer containing protease inhibitor cocktail and 20 mM NEM. Cell debris was removed by centrifugation, and the protein concentration in the supernatants was measured using the BCA Protein Assay kit (Thermo Fisher Scientific, Waltham, MA, USA). For immunoprecipitation, total cell lysates were incubated with anti-c-Maf (M-153/sc-7866; Santa Cruz Biotechnology, Dallas, TX, USA) overnight at 4 °C followed by incubation with 50 μL of protein G agarose (Millipore, Burlington, MA, USA) for 2 h at 4 °C. Immunoprecipitates were washed 5 times with RIPA buffer and subjected to SDS–PAGE and western blot analysis. After blotting, membranes were incubated overnight at 4 °C with anti-c-Maf (6B8/sc-293420; Santa Cruz Biotechnology) or anti-SUMO-1 (D-11/sc-5308; Santa Cruz Biotechnology) antibodies and subsequently incubated with goat anti-rabbit-conjugated HRP (Jackson ImmunoResearch Laboratories, West Grove, PA, USA) or goat anti-rat light chain-conjugate HRP (Jackson ImmunoResearch Laboratories) for 1 h at room temperature. In some cases, blots were incubated with Clean-Blot IP detection reagent (Thermo Fisher Scientific). Protein signals were detected using WesternBright ECL HRP substrate (Advansta Inc., San Jose, CA, USA) and visualized on an LAS-3000 imaging system (Fujifilm Life Science, Santa Ana, CA, USA).

### Full-length 16S rRNA gene sequencing

To characterize the gut microbiota composition, bacterial genomic DNA was extracted from fecal samples using the QIAamp Fast DNA Stool Mini Kit (Qiagen). The full-length 16S rRNA gene (V1–V9 regions) was amplified using barcoded primers (27F and 1492R), and libraries were sequenced on the PacBio Sequel IIe system (Pacific Biosciences, Menlo Park, CA, USA) to generate circular consensus sequencing reads. Data processing, including demultiplexing, quality filtering, and denoising (ASV inference), was performed using QIIME 2 and DADA2. Taxonomy was assigned against the SILVA database with a 99% identity threshold.

### Targeted metabolomics

To evaluate the functional metabolic output, fecal concentrations of SCFAs and BAs were quantified. For SCFAs, samples were acidified, extracted, and analyzed using gas chromatography-mass spectrometry on an Agilent 7890B system coupled to a 5977B MSD instrument (Agilent, Santa Clara, CA, USA). For bile acid profiling, specifically targeting LCA, DCA, and CDCA, lyophilized fecal extracts were analyzed using liquid chromatography-tandem mass spectrometry. Separation was performed on an ACQUITY UPLC system coupled to a Xevo TQ-S micro Triple Quadrupole Mass Spectrometer (Waters Corp., Milford, MA, USA) operating in multiple reaction monitoring mode.

### Generation of PBMC-humanized mouse model

The use of human biological materials in this study was approved by the Institutional Review Board of the Tri-Service General Hospital (TSGH; Taipei, Taiwan) (TSGHIRB No. E202516053). Cryopreserved human PBMCs from healthy donors were purchased from AllCells (Alameda, CA, USA). To establish the humanized mouse model, 4-week-old immunodeficient B6.*Rag2*^−/−^*Cd47*^−/−^*Il2rg*^−/−^ were used as recipients.

### Statistical analysis

The log-rank (Mantel–Cox) test was used to compare survival curves. The 2-tailed Student’s unpaired *t* test was used for comparisons between 2 groups. One-way ANOVA with Tukey’s multiple comparison test and 2-way ANOVA with Sidak’s multiple comparison test were used for multigroup comparisons. A *P* value <0.05 was considered to be significant.

### Study approval

All protocols using live animals were performed in accordance with institutional guidelines and were approved by the Institutional Animal Care and Use Committee at the National Defense Medical University (Taipei, Taiwan).

## Acknowledgments

We are grateful to the Health and Welfare Data Science Center (HWDC), Ministry of Health and Welfare (MOHW), Taiwan, for providing the National Health Insurance Research Database (NHIRD). We appreciate the technical support provided by BIOTOOLS Co., Ltd. for the 16S rRNA sequencing and targeted metabolomic analyses. We also thank the Core Instrument Center of National Defense Medical University for the assistance.

## Author contributions

CYH and YWT performed experiments and analyzed data; SHF, YWL, YWM, LCT, CEW and JLD performed experiments; YJY, HIL. CCS, CTC, SPW and SCM gave advice; CYH, YWT and HKS wrote the manuscript.

